# Dendritic Polyglycerol Amine Substrate Extends the Viability of Mixed Glial Cultures for Repeated Isolation of Immature Oligodendrocyte Lineage Cells

**DOI:** 10.1101/2025.06.23.661031

**Authors:** Nonthué A. Uccelli, Daryan Chitsaz, Jean-David Gothié, Divya Kakkar, Ehsan Mohammadifar, Jack P. Antel, Rainer Haag, Timothy E. Kennedy

## Abstract

Primary mixed glial cultures are key tools to isolate and study astrocytes, microglia and oligodendrocytes. Cell-substrate adhesion is critical for neural cell survival and differentiation. Cationic polymers like poly-D-lysine (PDL) are widely used to promote cell adhesion to cell culture substrates, however, PDL is not stable long-term, with cultured cells often detaching (peeling) after 2-3 weeks. Dendritic polyglycerol amine (dPGA) is a synthetic polycationic non-protein polymer biomimetic of poly-lysine that is highly resistant to degradation by cellular proteases. Substrates coated with dPGA promote cell adhesion and improve survival in long-term neuronal cultures. Here we assessed dPGA as a substrate coating to provide long-term support for mixed glial cultures. Oligodendrocyte precursor cells (OPCs) were isolated weekly by differential adhesion from cultures grown in T75 flasks with PDL or dPGA-coated substrates. Following two “shake-off” isolations, the cell layer in most PDL-coated flasks fully detached, rendering these flasks unusable for further culture. In contrast, dPGA-coated flasks consistently yielded cells for six or more sequential isolations over seven weeks in culture. dPGA-coated flasks produced more cells, a greater percentage of O4+ cells, and maintained similar proportions of OPCs and MBP-positive cells as when isolated from a PDL-coated substrate. dPGA is cyto-compatible, functionally superior, easy to use, low cost and a stable alternative to conventional cell substrate coatings. The enhanced long-term stability of mixed glial cultures grown on a dPGA substrate has the capacity to increase cellular yield, reduce animal use, and facilitate studies of oligodendrocyte cell biology.

## Introduction

Cell culture is an essential research tool that enables the study of cellular and molecular mechanisms under controlled conditions. Primary mixed glial cultures are widely used to obtain oligodendrocyte (OL) lineage cells, proliferating them in contact with an astrocyte monolayer, and isolating OPCs by differential adhesion using the shake-off technique (Armstrong, 1998; McCarthy and de Vellis, 1980). This method has been widely adopted by laboratories studying oligodendroglia cell biology. Here we extend the applicability of this technique to increase cellular yield, reduce the number of experimental animals required, and lower labor and reagent costs.

Primary cultures of adherent cells require a substrate to promote cell adhesion, survival, migration and process extension (Banker and Goslin, 1998). In early studies, purified extracellular matrix components were used to coat cultureware (Kleinman et al., 1987), however, it was found that substrates coated with synthetic positively charged amino acid polymers, like poly-lysine or poly-ornithine, significantly improved cell growth and survival (McKeehan and Ham, 1976). The use of synthetic polymers that are cationic at neutral pH avoids introducing impurities and variability associated with purified proteins or matrices, making them preferred alternatives to coat cell culture substrates (McKeehan and Ham, 1976; Yavin and Yavin, 1974). The gold standard substrate for mixed glial cell culture involves coating flasks with the polycationic polymer poly-D-lysine (PDL) (Armstrong, 1998; Banker and Goslin, 1998). Its enantiomer poly-L-lysine can also be used, but is less stable than PDL, as it is readily cleaved by secreted proteases (Banker and Goslin, 1998; Li and Yeung, 2008). Although PDL is resistant, it is not insensitive to cellular proteases. Degradation of the polymer or matrix underlying cells in culture results in cells “peeling” off the substrate, which limits mixed glial cultures to typically no more than two or three OPC isolations by “shake-off”.

Dendritic polyglycerol amines (dPGAs) are a family of hyperbranched polymers with a chemical structure composed of dendritic branching chains of glycerol monomers linked by ether bonds and terminal hydroxyl functional groups that are substituted with varying proportions of amines (Figure 1A) (Frey and Haag, 2002; Hellmund et al., 2015). Like PDL, dPGA carries a high density of positive charges at neutral pH, however, unlike the peptide bonds in PDL and PLL, the ether bonds in dPGA are not substrates for cellular proteases (Frey and Haag, 2002). Cell culture substrates coated with dPGA have been shown to support the adhesion and differentiation of rodent primary neurons and human iPSC derived neurons in long-term cell culture, with substantially increased stability compared to conventional PLL or PDL coatings (Clement et al., 2022; Thiry et al., 2022; Thiry et al., 2024).

**Figure 1.**
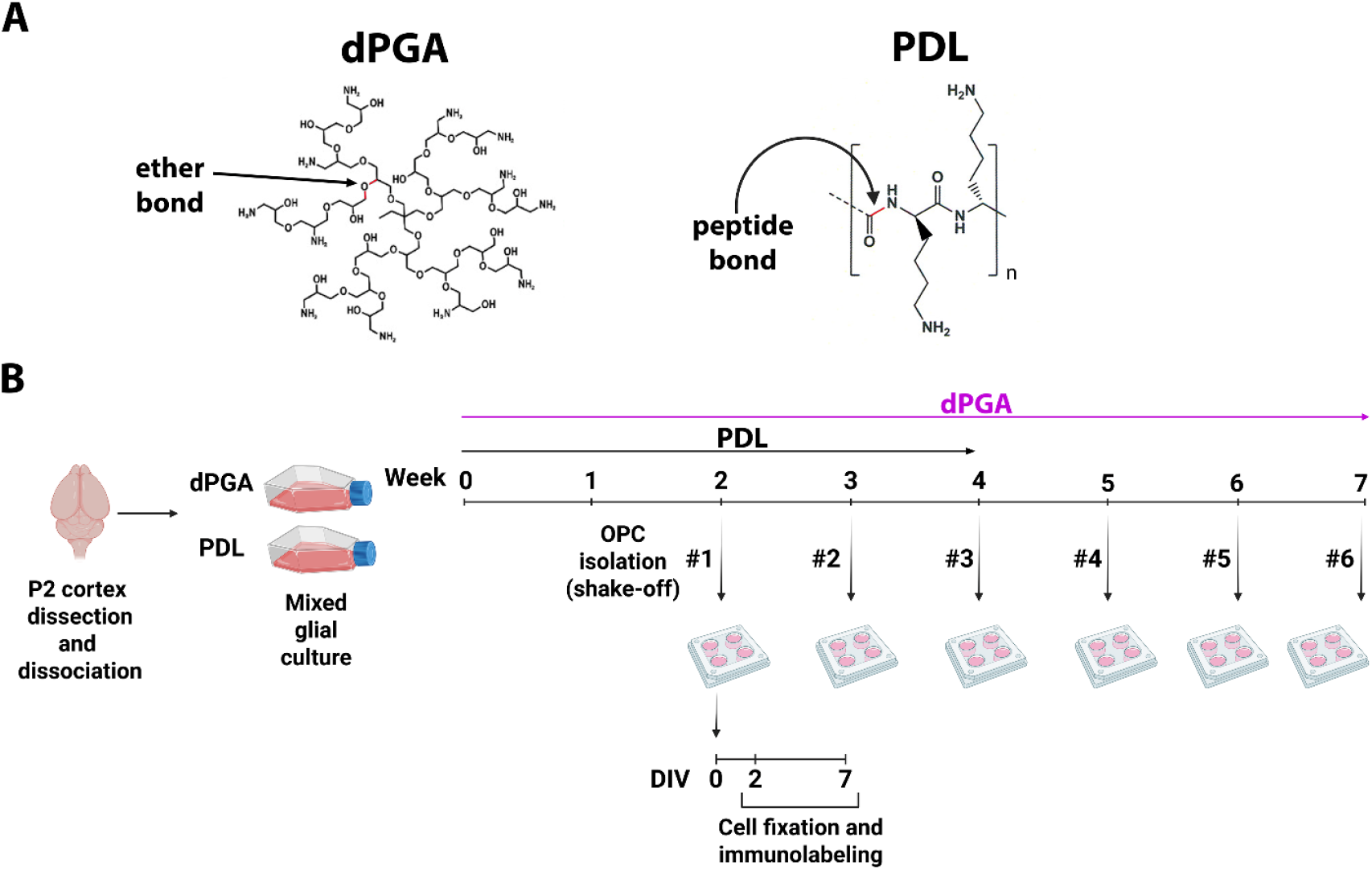
Chemical structures and experimental design. **A)** Illustrations of the chemical structures of dendritic polyglycerol amine (dPGA) and poly-D-lysine (PDL), highlighting their ether or peptide bonds, respectively. Adapted from Clement et al., 2022. **B)** Cortices from postnatal day (P2) rats were dissected and cells dissociated. The cell suspension was seeded into flasks with cell culture substrates coated with either dPGA or PDL and maintained undisturbed for two weeks. Beginning at the end of the second week, shake-off isolations were performed weekly, up to 7 weeks for the dPGA-coated flasks. After each isolation, cells were plated and maintained for either 2 or 7 days in vitro (DIV) before fixation and immunolabelling with oligodendrocyte lineage markers. Created with BioRender.

Here, we demonstrate enhanced performance of dPGA as a substrate coating for mixed glial cell cultures compared to PDL, increasing substrate stability, extending culture lifespan, and increasing the yield of cells obtained from each dissection.

## Materials and Methods

### Animals

Sprague Dawley rat pups were obtained from Charles River Canada (Montreal, Quebec, Canada). All procedures were performed in accordance with the Canadian Council on Animal Care guidelines for the use of animals in research and approved by the Montreal Neurological Institute Animal Care Committee (animal ethics protocol #4330).

### Cell culture

#### Substrate coating

The cell culture surfaces of T75 flasks (#83.3911.302, Sarstedt) were coated with PDL (10 ml of 10 μg/ml PDL per flask; 150–300 kDa, Sigma Aldrich #P1149) or dPGA (10 ml of 10 μg/ml dPGA per flask; ∼550 kDa, ∼30% amine functionalization, DendroTEK Biosciences, Montreal) diluted in PBS at pH 7.5. PDL-coated flasks were incubated at room temperature (rt) for 1 h, rinsed twice with dH_2_O, and dried before use (adapted from (Armstrong, 1998). dPGA-coated flasks were incubated at 37° C in an incubator with 5% CO_2_ for 1 h, and then rinsed three times with phosphate-buffered saline (PBS), without drying (adapted from (Clement et al., 2022). Each flask was filled with 9 ml of Dulbecco’s Modified Eagle Medium (DMEM, #11995073, Gibco) supplemented with 10% heat inactivated fetal bovine serum (FBS, # F1051, Sigma-Aldrich) and 1% filtered Penicillin/Streptomycin (#15140122, Gibco) (cDMEM) and kept in an incubator at 37°C with 5% CO_2_ (Forma series II, Thermo Scientific) for 30 min before adding the cell suspension.

#### Tissue dissection and dissociation

Primary mixed glial cultures were obtained by dissection of cerebral cortices from litters typically composed of ∼12 rat pups at postnatal day 2 (P2) (Figure 1B). In accordance with institutional guidelines, neonatal rats were rapidly decapitated, and brains isolated and placed in ice-cold HBSS. Isolation of the neocortex by dissection was performed in a petri dish with chilled 0.5 µg/ml Amphotericin B (#15290018, Gibco) in Hanks’ Balanced Salt Solution (HBSS, #14170161, Gibco) in a laminar flow hood workstation (Forma Scientific, model 1839).

For tissue dissociation, the dissected cortices were transported to a class II biosafety cabinet (1300 Series A2, Thermo Scientific) and finely chopped with an alcohol sterilized razor blade. The chopped tissue was incubated with 6 ml of 0.125% trypsin-EDTA (#25200056, Gibco) in HBSS for 20 min at 37°C, adding DNase I (#DN25, Sigma-Aldrich) in HBSS with calcium and magnesium chloride (#24020117, Gibco) at a concentration of 1 mg/ml for the last 5 min. After removing tissue from the incubator, it was resuspended in a 50 ml tube and centrifuged for 5 min at 241 x g (IEC Centra CL2). After aspirating the supernatant, the pellet was resuspended in 10 ml of cDMEM and mechanically dissociated. After passing through a 70 µm pore cell strainer (#22-363-548, Fisherbrand), 1 ml of cell suspension was added to each flask, resulting in each flask containing dissociated cortices from ∼1.2 pups. For each dissection performed in this study (n=5), half of the cells obtained were seeded in PDL-coated flasks and the other half in dPGA-coated flasks. All flasks were kept in the incubator at 37 °C in 5% CO_2_ for 2 weeks before the first isolation (Figure 1B).

#### OPC isolation from mixed glial cell cultures by differential adhesion

A pool of cells enriched for oligodendrocyte precursor cells (OPCs) was isolated by differential adhesion following a mechanical shake-off protocol, essentially as described (Armstrong, 1998; Jarjour et al., 2003). Flasks were first sealed with Parafilm (#52858-000, Bemis) and shaken for 1 h at 37°C, at 150 rpm in an orbital shaking incubator with a rotational diameter of 3 cm (Jeio Tech, model SI600), to detach the microglia, but not the OPCs or astrocytes, which remain attached to the bottom of the flask. Detached microglia were removed with the culture medium. Fresh cell culture medium was then added to each flask, and the cultures allowed to recover for 1 h at 37 °C in 5% CO_2_ without shaking. The flasks were then returned to the shaking incubator, and shaken again, at 37 °C and 180 rpm overnight. This more intense shaking aims to detach the OPCs, while leaving the underlying layer of astrocytes intact. The following day, the detached OPCs suspended in the cell culture medium were collected from each flask, transferred to a standard polystyrene petri dish (#82.1473.001, Sarstedt) and then kept in the cell culture incubator for 20 min. This step aims to remove any astrocytes that may have detached from the flask by preferential adherence to the bottom of the petri dish while the OPCs remain in suspension. The cell culture medium containing the OPCs was then collected and transferred to 50 ml tubes and centrifuged for 7 min at 168 x g. The resulting OPC enriched cell pellets were resuspended in SATO medium (DMEM supplemented with 5 μg/ml insulin, 50 μg/ml transferrin, 30 nM sodium selenite, 30 nM triiodothyronine, 6.3 ng/ml progesterone, 16 μg/ml putrescine, 0.5% FBS and 2 mM glutamax), mechanically triturated with a 21-gauge needle and transferred through a cell strainer. Cells were counted with a hemocytometer and the yield calculated (#cells per flask). OPC isolations were performed from the same flasks, once a week.

#### Substrate coating and OPCs seeding

Individual round cover glasses (12 mm diameter, No. 0, #92100100030, Carolina) were placed into each well of a 24-well cell culture plate and sterilized with a plasma cleaner for 1 min (Harrick Plasma). They were then coated with 10 μg/ml in PDL for 1 h at rt, then rinsed twice with dH_2_O and allowed to dry. Isolated OPCs and oligodendrocytes were seeded onto PDL-coated cover glasses at a density of 50,000 cells/coverglass and maintained in SATO medium at 37°C in 5% CO_2_ for 2 or 7 days *in vitro* (DIV) for OPC and mature OL enriched cultures, respectively.

### Immunocytochemistry

Mixed glial cultures supported by substrates coated with PDL or dPGA were fixed in the flasks 14 days after dissection with 4% paraformaldehyde (PFA) and washed with PBS. The plastic base of the T75 flasks with cells attached was cut out with a metal micro-spatula heated over a Bunsen burner and then cut into 2 cm^2^ pieces which were placed in a 6-well dish for immunolabelling. Antibodies against neural/glial antigen 2 (NG2, 1:500, Milipore #MAB5384), ionized calcium binding adaptor molecular 1 (Iba1, 1:300, Wako #019-19741), and glial fibrillary acidic protein (GFAP, 1:1000, Milipore #AB5541) were used to label OPCs, microglia, and astrocytes, respectively, along with Hoechst dye (#33342, Thermo Fisher Scientific, Montreal) to label nuclei. The immunolabeled pieces of the flask base were then mounted with Fluoro-Gel (Electron Microscopy Sciences), coverslipped, and sealed with UV-curing nail polish (Kennedy et al., 2023).

Following 2 or 7 DIV after isolation, cells were fixed for 10 min with 4% PFA and washed with PBS. Cells were blocked with PBS containing 2.5% bovine serum albumin and 2.5% horse serum with 0.05% Triton X-100 for 30 min at rt. Cells were incubated overnight at 4°C with primary antibodies against myelin basic protein (MBP, 1:1000; Aves #MBP0020), platelet-derived growth factor alpha receptor (PDGFαR, 1:500; C-20, Santa Cruz Biotechnology) and oligodendrocyte marker O4 (1:500, R&D Systems #MAB1326) diluted in blocking solution. Following primary antibody washes, cells were labeled with fluorescent secondary antibodies Alexa Fluor 488, 555, 647 (1:1000; Invitrogen) diluted in PBS with 2.5% bovine serum albumin and 2.5% horse serum, with 5 µg/ml Hoechst dye, and incubated for 1 h at rt. Two washes with PBS were performed before mounting the coverslips with Dako mounting media.

### Imaging

To visualize mixed glial cell cultures on PDL and dPGA substrates, immunolabeled flask substrates were imaged using a Zeiss LSM-880 Airyscan 1 confocal microscope with a 20x objective (n.a. = 0.8) (Carl Zeiss Canada, Toronto). Live cells in flasks were imaged before each shake-off using a 20x Variable Relief (VAREL) contrast objective (n.a. = 0.3, LD A-Plan, Zeiss) on a Zeiss Axiovert S100TV inverted microscope. The detaching layer of astrocytes was imagined using DIC with a 10x objective (n.a. = 0.45, Plan-Apochromat, Zeiss) on a Zeiss Axio Observer.Z1 microscope equipped with an Axiocam 506 camera.

To analyse the composition of the oligodendroglial population after 2 or 7 DIV post-isolation, regions of interest on each cover glass were selected randomly and imaged with a 20x objective (n.a. = 0.8) using a Zeiss Axio Observer.Z1 microscope equipped with an Axiocam 506 camera. MBP, PDGFαR and O4 immunopositive cells were counted using the ImageJ Cell Counter plugin (Schneider et al., 2012). The number of DAPI positive cells was counted using a semi-automated threshold and mask strategy in ImageJ.

### Statistics

Statistical analyses were performed using GraphPad Prism 8.0 (GraphPad Software, Boston, MA). Shake-off yields were compared by two-way analysis of variance (ANOVA) followed by Sidak’s multiple comparison test. After assessing normality and homogeneity of variance, the number or fraction of cells were compared by parametric two-way analysis of variance (ANOVA) followed by Tukey’s multiple comparison test or by non-parametric Kruskal-Wallis followed by Dunn’s multiple comparison test.

## Results

### dPGA substrate enhances astrocyte monolayer stability to extend mixed glial culture lifetime

Here, we investigated the efficacy of coating substrates with dPGA to support long-term mixed glial cell cultures. The neocortex was dissected from P2 rats, cells dissociated, and then plated in T75 flasks with cell culture substrates coated with dPGA or PDL. After 2 weeks, these cultures typically establish a relatively uniform layer of astrocytes that cover the cell culture surface of the flask, with microglia and OPCs growing on top of the astrocyte base layer.

Following 2 weeks, the largely confluent layer of cells was fixed in the T75 flasks and immunolabeled for NG2 to identify OPCs (Nishiyama et al., 1999), Iba1 to mark microglia (Ito et al., 1998), and GFAP to identify astrocytes (Eng et al., 2000; Eng et al., 1971) (Figure 2A). Overall, the layer of cells, before the first shake-off, appeared similar on either PDL or dPGA substrates, yet more gaps in the cell layer were visible in the PDL-coated flasks with more complete coverage of the surface in the dPGA-coated flasks. The small round phase-bright cell bodies of OPCs were readily visible on top of the astrocyte layer before the first shake-off from either coated surface (Figure 2A).

**Figure 2.**
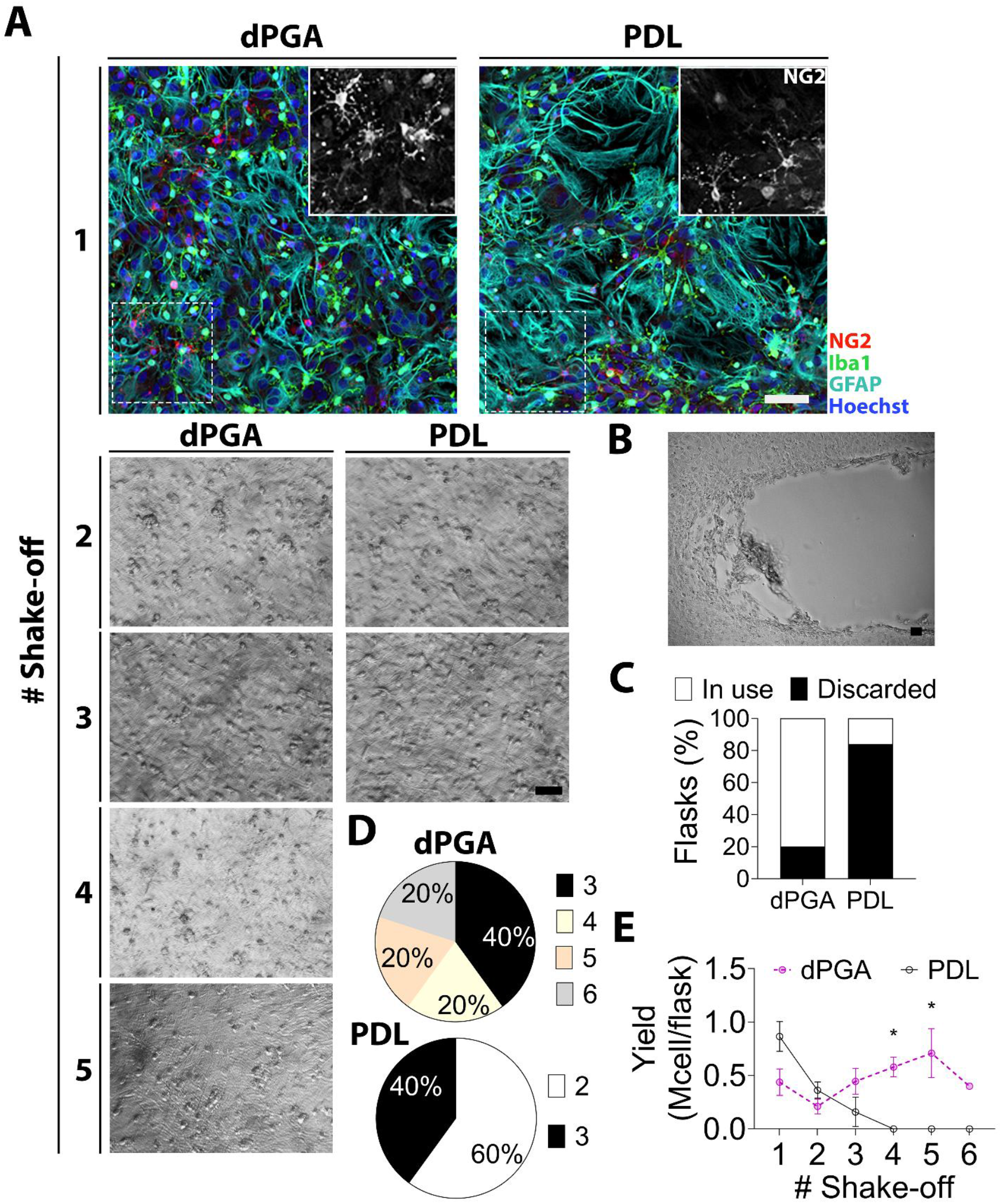
dPGA extends the viability of mixed glial cultures, enabling multiple cell isolations. **A)** Images of mixed glial cell cultures on dPGA or PDL coated substrates, imaged before subsequent shake-offs (1-5). Following 2 weeks without a shake-off, cells were fixed within the flasks and then immunolabeled for glial markers to identify OPCs (NG2), astrocytes (GFAP) and microglia (Iba1). For subsequent shake-offs, live cells were imaged in the flasks using Variable Relief (VAREL, 20x objective) contrast microscopy. **B)** Micrograph shows the edge of an astrocyte monolayer detaching from a PDL coated substrate. After two to three shake-offs from a PDL coated substrate the astrocyte monolayer typically begins to peel and detach, in which case the flasks must be discarded. **C)** Following a 3^rd^ shake-off, most PDL-coated flasks were discarded, while 80% of dPGA-coated flasks remained viable for further OPC isolations. **D)** Across all dissections (n=5), dPGA-coated flasks provided 3-6 or more shake-offs, compared to 2-3 shake-offs for PDL-coated flasks. **E)** Number of cells isolated per flask on each shake-off from PDL verses dPGA-coated flasks. All scale bars = 50 µm.

To generate a pool of cells enriched with OPCs, shake-off isolations were performed, based on differential adhesion of the different glial cell types to different substrates (Armstrong, 1998). Following this technique, a population enriched with microglia is first removed from the T75 flask with a relatively brief and gentle shake (1 h at 150 rpm), while the OPC enriched pool is detached from the astrocyte layer by shaking at a higher speed overnight. Images of live mixed glial cell cultures growing in the flasks were obtained using VAREL contrast imaging before each isolation (performed weekly), to visualize the integrity of the cell layer on the different substrate coatings (Figure 2A). Using PDL coated substrates, following the second to third shake-off, the base layer of astrocytes began to peel away from the substrate (Figure 2B), with approximately 80% of the cultures on PDL coated substrates completely detached by the third shake-off (Figure 2C). In contrast, approximately 80% of the cultures maintained on dPGA-coated substrates remained intact (Figure 2C, D). Following the first and second isolations, comparable numbers of cells were obtained from the PDL and dPGA-coated flasks (Figure 2E). In our hands, none of the PDL coated flasks were usable after a third shake-off (Figure 2D). In contrast, on dPGA-coated substrates, the layer of astrocytes remained intact, allowing further rounds of cell isolation to be performed, maintaining yields of ∼0.5 million cells/flask for up to 7 weeks (Figure 2D-E).

### Increased numbers of OPCs isolated from dPGA substrates without altering the proportion of mature oligodendrocytes

We then further characterized the isolated cells to determine if maintaining a mixed glial culture on a dPGA-coated substrate might alter the population of oligodendrocyte lineage cells obtained after isolation relative to cells cultured on a PDL substrate. To provide a direct comparison of the cells derived from mixed glial cultures supported by dPGA verses PDL substrates, all isolated OPCs were replated onto coverslips coated with PDL. Following replating, cells were fixed and immunolabeled after 2 and 7 DIV to identify OPCs (PDGFαR), immature and early mature oligodendrocytes (O4), and mature myelinating oligodendrocytes (myelin basic protein, MBP) (Figure 3). For quantification and comparison of the PDL verses dPGA coated surfaces, the values obtained for cells isolated from the first and second shake-offs were combined (PDL_1/2_, dPGA_1/2_), and compared with the combined values of cells isolated from dPGA-coated flasks from the third and fourth shake-off (dPGA_3/4_).

**Figure 3.**
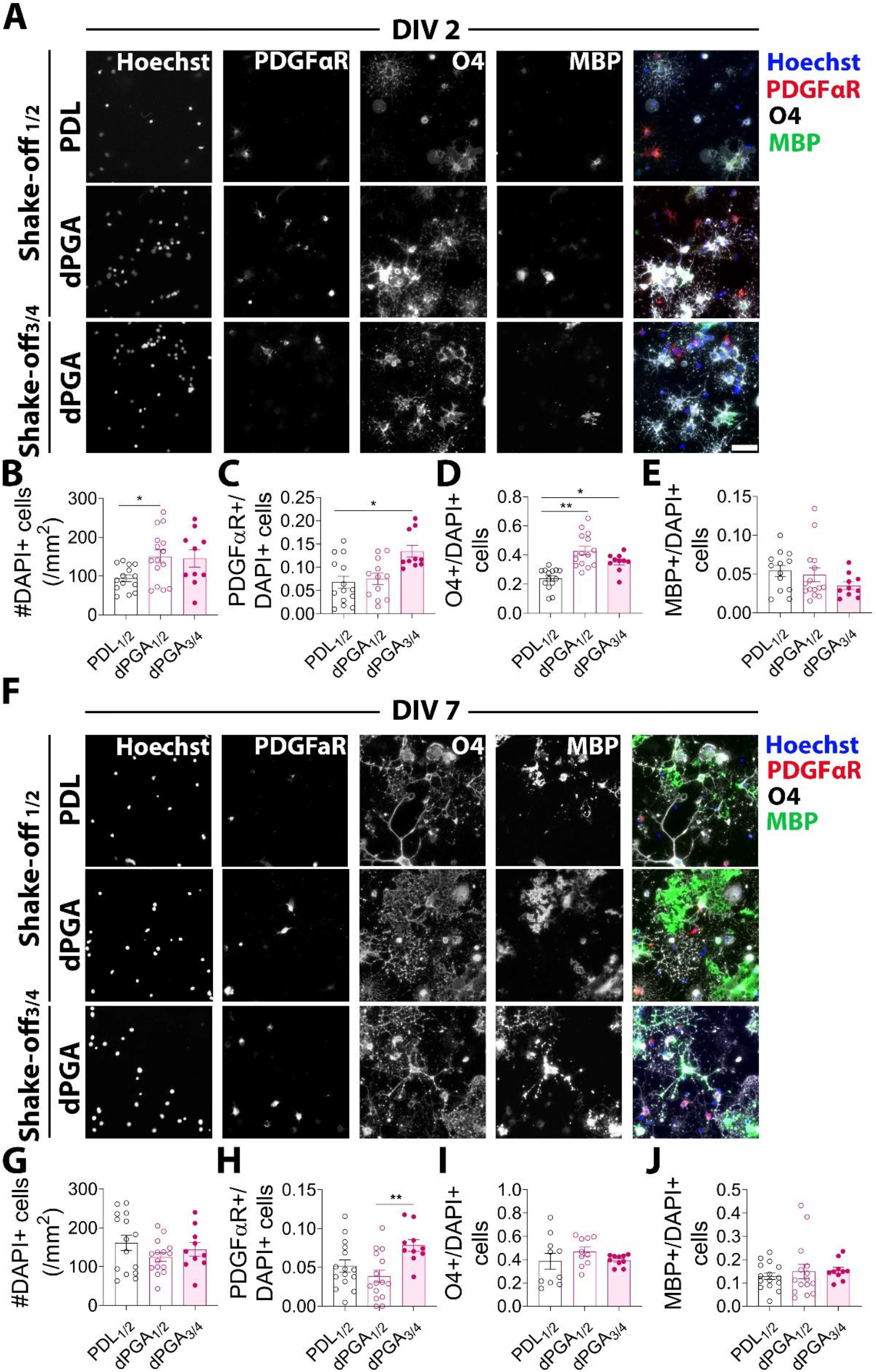
Lineage profile of OPCs and oligodendrocytes isolated from dPGA verses PDL-coated flasks comparing early verses late isolations. OPCs were isolated from mixed glial cell cultures by sequential shake-offs. dPGA or PDL-coated substrates supported at least 2 isolations (PDL_1/2_, dPGA_1/2_), however, a dPGA coating provided a more stable substrate that supported longer lasting cultures, typically allowing 3-6 or more additional isolations. To evaluate OPC cultures obtained in later isolations, cells were pooled from the 3^rd^ and 4^th^ shake-off (dPGA_3/4_). **A)** OPCs were isolated from dPGA or PDL coated flasks, replated, and immunolabeled following 2 DIV to identify OPCs (PDGFαR positive), immature/early mature oligodendrocytes (O4 positive) and mature myelinating oligodendrocytes (MBP positive). **B)** dPGA-coated substrates supported a higher density of cells (DAPI positive) than PDL coated substrates. **C)** Relative to the total number of cells isolated, similar fractions of PDGFαR-positive cells were obtained from early isolations derived from dPGA or PDL-coated flasks. Later isolations from dPGA coated flasks (dPGA_3/4_), produced a significantly larger proportion OPCs compared to the earlier isolations. **D)** A larger proportion of O4-positive cells were obtained when isolated from dPGA-coated substrates compared to PDL-coated substrates. **E)** Similar proportions of MBP-positive cells were obtained regardless of substrate coating or culture length. **F)** After isolation from dPGA or PDL coated flasks in early (PDL_1/2_, dPGA_1/2_) or late (dPGA_3/4_) shake-offs, OPCs were fixed at 7 DIV and then immunolabeled for oligodendrocyte lineage markers: PDGFαR, O4 and MBP. **G)** At 7 DIV, cell density (DAPI positive) was similar across groups. **H)** After 7 DIV, cells isolated from PDL and dPGA-coated substrates exhibited similar numbers of OPCs, but older cultures (dPGA_3/4_) presented increased numbers of PDGFαR-positive cells. **I)** The fraction of O4-positive oligodendrocytes was similar across groups at 7 DIV after isolation. **J)** MBP-positive cell numbers were similar across all groups at 7 DIV after isolation. Scale bars= 50µm.

At 2 DIV after replating on PDL there were approximately 30% more cells (DAPI-positive) in cultures derived from mixed glial cultures grown on a dPGA-substrate compared to cultures derived from PDL-coated flasks (Figure 3B). Further, close to a doubling of the proportion of O4-positive oligodendrocytes was obtained for cells derived from dPGA-coated flasks (Figure 3D). Considering that the same total number of cells were initially replated, these findings indicate that the cells derived from dPGA supported mixed glial cultures are more proliferative and enriched with oligodendrocytes. The fraction of PDGFαR immunopositive OPCs did not differ between PDL or dPGA groups isolated in the first or second shake-offs, however, later isolations from dPGA coated flasks produced both more cells (Figure 3B) and a larger proportion of PDGFαR immunopositive OPCs (Figure 3C). The fraction of MBP-positive cells was similar across groups and accounted for a small portion of the replated cells, as expected after only 2 DIV (Figure 3E).

After 7 DIV on PDL coated coverslips, many of the OPCs obtained from mixed glial cultures following this protocol have differentiated into oligodendrocytes (Armstrong, 1998; McCarthy & de Vellis 1980). To determine if maintaining the mixed glial cultures on a dPGA substrate might influence subsequent oligodendrocyte differentiation, we then characterized the population of oligodendrocytes present at 7 DIV following replating. Comparing the cells derived from the first and second shake-off isolations from dPGA verses PDL coated flasks, similar total numbers and proportions of PDGFαR immunopositive OPCs, and O4 and MBP-positive oligodendrocytes were detected one week following replating, irrespective of the substrate coating used for the mixed glial cultures (Figure 3 G-J). Isolation from more mature mixed glial cultures, following three to four shake-offs from dPGA substrates, produced a larger fraction of PDGFαR-immunopositive OPCs, detected at 2 and 7 DIV following replating (Figure 3 C, H). These results provide evidence that a dPGA substrate enhances the generation of OPCs in mixed glial cell culture, and that OPCs generated in mixed glial cell cultures supported by dPGA or PDL substrates have similar capacity to differentiate to mature oligodendrocytes.

## Discussion

We present dPGA as a novel substrate coating to support enhanced long-term stability of mixed glial cell cultures. We show that cultures grown on dPGA-coated substrates are comparable to those on PDL-coated substrates in terms of cellular composition, however, the stability of the dPGA coating allows the cultures to be maintained for substantially longer periods of time, supporting additional cycles of cell isolation using the shake-off protocol and increasing the yield of OPCs generated.

Both PDL and dPGA are large polymers that present positively charged amines at neutral pH which act as a bridge between the negative surface charge of the cell culture substrate and the negative charge presented by plasma membrane phospholipids and glycosaminoglycans (Calderon et al., 2010; Clement et al., 2022; Pouyan et al., 2022). The binding mediated by both cationic polymers likely engages similar mechanisms of cell-substrate adhesion and differentiation. Consistent with this, following shake-off isolation and replating, we show that by 7 DIV the proportion of differentiated oligodendrocytes generated from OPCs is similar, regardless of whether the mixed glial cultures were maintained on PDL or dPGA-coated substrates.

By 2 DIV, cultures derived from dPGA-coated flasks and replated exhibit an increase in total cell number and a higher proportion of O4-immunopositive oligodendrocytes compared to cultures derived from PDL-coated flasks. Since the proportions of OPCs and mature oligodendrocytes remain unchanged, this suggests that mixed glial cultures supported with a dPGA-coated substrate yield a greater number of oligodendrocyte lineage cells. Employing additional steps of purification or selectively promoting OPC proliferation using mitogens could be used to further increase the enrichment and yield of oligodendrocytes in these cultures.

Substrates coated with dPGA, compared to PDL, better support the long-term stability of mixed glial cultures, allowing for additional isolations of primary cells from each dissection. Of interest, prolonged culture in dPGA-coated flasks resulted in an increase in the fraction of OPCs isolated from later shake-offs. The increased stability of the mixed glial culture, in particular the enhanced support provided for the underlying layer of astrocytes, with reduced cell clumping and peeling, likely contributes to the proliferation of additional OPCs. As described for human iPSC derived motor neurons cultured on a dPGA coated substrate (Thiry et al., 2022; Thiry et al., 2024), the extended stability and longevity of mixed glial cultures supported by dPGA may facilitate the study of differences that emerge in more mature cells (Heo et al., 2024; Nishiyama et al., 2021; Sim et al., 2002) and provide opportunities to generate cells *in vitro* that better reflect phenotypes associated with adult neural degenerative diseases..

A key advantage of dPGA over poly-lysine and other protein-based matrices is its superior stability. Critically, dPGA is not susceptible to degradation by cellular proteases that target peptide bonds. Future studies could explore how variations in the properties of dPGA, such as polymer size and charge might be tuned to further enhance the stability of the astrocyte monolayer and promote the proliferation of OPCs. Such improvements could further extend the lifespan of mixed glial cultures and expand their utility in experimental applications.

The method described here represents a substantial improvement in yield of oligodendrocyte lineage cells compared to mixed glial cultures grown on a conventional poly-lysine coated substrate. A dPGA substrate coating enables more oligodendroglial cells to be obtained over an extended number of cycles of cell isolations using the differential adhesion shake-off technique. This enables more efficient use of resources, with increased numbers of isolations possible from a single dissection reducing the time, reagents and cost required to generate OPCs. This is cost effective, promotes laboratory sustainability (Freese et al., 2024), and reduces the number of animals required for studies using primary oligodendrocytes (CCAC, 2017; Russell and Burch, 1959).

## Acknowledgments

The project was supported by grants from the Multiple Sclerosis Society of Canada -(1038154, TEK), NSERC-CREATE - (575235-2023, TEK), and IRTG 2662 funded by the Deutsche Forschungsgemeinschaft (DFG, German Research Foundation) - (#434130070, RH). DC was supported by fellowships from the Multiple Sclerosis Society of Canada and the Fonds de la Recherche en Santé du Quebec, and NAU by Hoffmann-La Roche Canada, McGill MNI Jeanne-Timmins Costello and Redpoll post-doctoral fellowships.

